# Worms of change: anthropogenic disturbance changed the ectoparasite community structure of Lake Victoria cichlids

**DOI:** 10.1101/2024.04.14.589059

**Authors:** Tiziana Gobbin, Maarten Van Steenberge, Nathan Vranken, Maarten PM Vanhove

**Affiliations:** Research Group Zoology: Biodiversity & Toxicology, Centre for Environmental Sciences, Hasselt University, Diepenbeek, Belgium; Operational Directorate Taxonomy and Phylogeny, Royal Belgian Institute of Natural Sciences, Brussels, Belgium; Laboratory of Biodiversity and Evolutionary Genomics, Department of Biology, University of Leuven, Leuven, Belgium; Royal Museum for Central Africa, Biology department, Section Vertebrates, Tervuren, Belgium

**Keywords:** host-parasite interactions, anthropogenic disturbance, parasite conservation, global change

## Abstract

Host-parasite interactions increase the complexity, and thus robustness and resilience, of an ecosystem. This role is particularly relevant in times of global change. Environmental changes cause biodiversity loss and shifts in community compositions of free-living organisms, but how these changes affect parasite communities is still unclear. We tested how parasites respond to anthropogenic perturbations, using the case of Lake Victoria (East Africa), after 40 years of their onset. Lake Victoria experienced multiple human-induced invasions (e.g. Nile perch) and eutrophication that heavily affected haplochromine cichlid fishes (whose species richness decreased from 500 to 250 species in a decade). We compared gill macroparasite communities of 13 haplochromine species before and after perturbations, using historical and recent fish collections. The host-parasite network re-arranged, with few emerging and few disappearing host-parasite combinations, in a way that reduced stability. The host range of parasites, which is linked to resilience ability, decreased and thus we expect a decreased resilience ability in the future. We also found a decrease in infection parameters, co-infection occurrence, and biodiversity indices highlighting the urgent need of a conservation plan for wildlife parasites, to preserve their ecosystem services in face of global change. This study serves as a proof-of-concept of how often-overlooked aspects of host-parasite interactions provide a tool to monitor the health status of an ecosystem.

## 1. Introduction

A more complex ecosystem is more robust to change(s) than a simple one (Pimm 1984, Landi et al. 2018). Host-parasite interactions increase ecosystem complexity and thereby its robustness (i.e. ability to maintain the original status after perturbations) and resilience (i.e. ability to return to a previous state or establish a new state after perturbations) (Lafferty et al. 2006, Lafferty et al. 2008, Dunne et al. 2013), which is particularly valuable in the current global change era. Parasite abundance and diversity are expected to change in response to anthropogenic perturbations, however, the direction of that change is still object of debate (some studies found that parasitism was favored, others that it was disadvantaged, Blanar et al. 2009, Scharsack et al. 2021), let alone its consequences for ecosystem robustness and resilience.

For parasites, environmental changes encompass changes in host ecology and/or communities (as hosts represent an environment for parasites) along with abiotic changes (e.g. drought, temperature rise, pollutants). Ecosystem changes may lead to decline or extinction in native host species, potentially causing parasite (co-)extinction (Mey 1990, Faust et al. 2018, Šimková et al. 2019), parasite rising in abundance (Scharsack et al. 2021), parasites switching to new host species (Roche et al. 2010, Kafle 2018, Jorissen et al. 2020), shifts in host range (Ricklefs et al. 2004, Weckstein 2004) and even zoonotic spillovers (Jones et al. 2013, Allen et al. 2017, Faust et al. 2018, Johnson et al. 2020). These events can constitute a **conservation risk** for parasites (co-extinction) and a **disease outbreak risk** for hosts (changes in host use), including humans (spillovers). Knowing how parasite communities are varying across time can indicate how ecosystem complexity changed, even before we could observe changes in free-living organisms. Thus, changes in parasite communities can inform us on the health status of an ecosystem (**indicator potential**) (Shea et al. 2012, Hernández-Camacho and Zamora-Ledesma 2023).

Long-term parasite data can shed light on the direction of change in parasitism, but these are lacking, as the average dataset spans over 12 years (Fiorenza et al. 2020b). Parasitological dissections of specimens from natural history collections can provide the necessary time coverage to reconstruct changes in parasite abundance and diversity in response to ecosystem degradation, and allows for community ecological studies because each host specimen harbours a parasite community.

We test how parasite infection and host-parasite interactions change in response to ecosystem perturbations, using the case of Lake Victoria (eastern Africa). Lake Victoria is a biodiversity hotspot, harbouring a young and fast adaptive radiation of ~500 haplochromine cichlids (Cichlidae: Haplochromini) that evolved *in situ* in the past 14,600-16,000 years (Meier et al. 2017, McGee et al. 2020, Wienhues et al. 2024). Since the 1950s, human activities heavily impacted Lake Victoria and, often with a cascade effect, these resulted in drastic ecosystem changes. The exotic Nile perch (*Lates niloticus*) was introduced in northern Lake Victoria in the 1950s-1960s (Pringle 2005, Goudswaard et al. 2008) and became an **invasive species** in the 1980s. In Lake Victoria, the Nile perch switched diet to haplochromine cichlid fishes, the locally most abundant prey (Gee 1969), negatively impacting their populations. Concomitantly, **eutrophication** (i.e. water enrichment in inorganic nutrients) increased in Lake Victoria, due to urban/agricultural pollution and soil erosion through deforestation (Verschuren et al. 2002). This altered phytoplankton composition (Hecky 1993, Verschuren et al. 2002), increased water turbidity (Seehausen et al. 1997), reduced dissolved oxygen hence expanding deep anoxic zones (Wanink et al. 2001) and caused algal blooms (Witte et al. 1992a, Sitoki et al. 2010). These anthropogenic factors led to shifts in community compositions (e.g. fish, Witte et al. 2007a, Witte et al. 2007b; phytoplankton, Sitoki et al. 2012; zooplankton, Gophen et al. 1995) and biomass (increased in phytoplankton Wakwabi et al. 2006; decreased in benthic macroinvertebrates, Ngupula and Kayanda 2010) and an overall rapid and drastic **biodiversity loss** (Witte et al. 1992b, Seehausen et al. 1997, Witte et al. 2007a). Haplochromine cichlid fishes were particularly affected, as their decline is estimated from 500 to 250 species in a few decades (Witte et al. 1992b). Niches previously occupied by haplochromines were taken over by other species, making Lake Victoria dominated by non-haplochromine fishes, simplifying considerably the food web (Witte et al. 1992a, Witte et al. 1992b).

We screened 13 haplochromine species from recent and historical fish collections for ectoparasites to test whether and how parasite communities changed in response to (anthropogenic) ecosystem perturbations in Lake Victoria and whether parasites can be used as indicators of changing ecosystem health. Because of reduced abundance and diversity of host haplochromines, we expect a reduction in the abundance of their parasites, especially for those with high host specificity, and a change in host range.

## 2. Materials and methods

### 2.1. Fish collection

Cichlid fishes (n=340) were obtained from the ichthyological collections of the Naturalis Biodiversity Center (Leiden, the Netherlands) and of the Swiss Federal Institute of Aquatic Science and Technology (EAWAG, Kastanienbaum, Switzerland). These specimens were originally sampled during different field campaigns at different locations in Lake Victoria between 1973 and 2014. In particular, most fish were collected during an expedition of the ‘*Haplochromis* Ecological Survey Team’ in 1978-9 (HEST 1988) and the Bern University-EAWAG campaigns in 2010 and 2014 (Karvonen et al. 2018, Gobbin et al. 2020, Gobbin et al. 2023). Both took place in Speke Gulf and Mwanza Gulf (southern part of the lake). Most fish (n=275) were preserved in 4% formalin and subsequently transferred to 70% ethanol; other fish (n=65) were directly preserved in 100% ethanol for future genetic analysis.

We selected 13 common haplochromine species (nomenclature following Froese and Pauly 2000) belonging to 7 trophic groups representing the trophic diversity of Lake Victoria haplochromines (based on teeth morphology and/or stomach content, Seehausen 1996, Seehausen and Bouton 1998; **Table 1**), with specimens available from both before and after ecosystem perturbations. We set 1980 as the temporal cut-off, because in that year the Nile perch became invasive also in southern Lake Victoria and water eutrophication became apparent (Ochumba and Kibaara 1989, Sitoki et al. 2010). All haplochromine species sampled belong to the Lake Victoria radiation lineage, except *Astatoreochromis alluaudi* that represents an older and distantly related lineage that never radiated in Lake Victoria. We screened at least 10 fish individuals per species and per time period (before vs. after perturbations) for gill macroparasites. Each fish individual was also measured (standard length, to the nearest mm) and sexed.

**Table 1 -.** Host species of Haplochromis (Cichlidae: Haplochromini) sampled before and after perturbations in Lake Victoria. N = number of fish individuals; SL = standard length (mm), indicated as mean and range (minimum-maximum).

|  | Host species | Trophic group | N | SL (mm) |  |
| --- | --- | --- | --- | --- | --- |
|  |  |  |  | mean | (min-max) |
| before<br>perturbations | *Haplochromis mbipi* | algivore epilithic | 10 | 76.9 | (66.5-83.3) |
|  | *Haplochromis obliquidens* | algivore epiphitic | 10 | 72.7 | (67.8-78.2) |
|  | *Haplochromis antleter* | detritivore | 11 | 64.4 | (59.8-70.2) |
|  | *Haplochromis chilotes* | insectivore | 11 | 93.1 | (70.3-110.3) |
|  | *Haplochromis riponians* | insectivore | 10 | 104.3 | (91.4-112.9) |
|  | *Haplochromis sauvagei* | insectivore | 10 | 74.6 | (57.7-83.4) |
|  | *Haplochromis* sp. 'yellow chin pseudonigricans' | insectivore | 10 | 79.5 | (69.5-86.0) |
|  | *Astatoreochromis alluaudi* | molluscivore | 18 | 70.8 | (48.9-129.0) |
|  | *Haplochromis ishmaeli* | molluscivore | 10 | 107.6 | (94.8-121.5) |
|  | *Haplochromis xenognathus* | molluscivore | 10 | 78.9 | (59.0-100.5) |
|  | *Haplochromis spekii* | piscivore | 11 | 176 | (144.9-191.4) |
|  | *Haplochromis nyererei* | planktivore | 11 | 61.9 | (50.0-70.0) |
|  | *Haplochromis pyrrhocephalus* | zooplanktivore | 10 | 71.4 | (66.0-78.8) |
| after<br>perturbations | *Haplochromis mbipi* | algivore epilithic | 20 | 92.5 | (73.9-113.2) |
|  | *Haplochromis obliquidens* | algivore epiphitic | 10 | 68.3 | (59.7-104.1) |
|  | *Haplochromis antleter* | detritivore | 10 | 55.2 | (47.6-61.7) |
|  | *Haplochromis chilotes* | insectivore | 20 | 102.4 | (65.9-126.5) |
|  | *Haplochromis riponians* | insectivore | 10 | 90.2 | (76.6-101.5) |
|  | *Haplochromis sauvagei* | insectivore | 22 | 102.8 | (88.0-115.4) |
|  | *Haplochromis* sp. 'yellow chin pseudonigricans' | insectivore | 10 | 92.1 | (79.7-113.0) |
|  | *Astatoreochromis alluaudi* | molluscivore | 21 | 94.3 | (48.2-133.7) |
|  | *Haplochromis ishmaeli* | molluscivore | 15 | 79.1 | (0.0-112.9) |
|  | *Haplochromis xenognathus* | molluscivore | 11 | 105.3 | (80.8-115.4) |
|  | *Haplochromis spekii* | piscivore | 4 | 145.3 | (130.8-170.7) |
|  | *Haplochromis nyererei* | planktivore | 15 | 83.9 | (74.2-106.7) |
|  | *Haplochromis pyrrhocephalus* | zooplanktivore | 30 | 49.9 | (41.7-58.7) |

We use a novel approach: historical museum collections to reconstruct changes in gill parasite assemblages infecting fish (Wood et al. 2023a, Wood and Vanhove 2023). The advantages of this approach are: *i)* each host individual harbors a parasite community, allowing community analysis, *ii)* parasite detectability is not influenced by museum preservation protocols (Fiorenza et al. 2020a), and *iii)* gill parasites are physically protected by the fish operculum thus infection levels are likely unaltered during host manipulation (Kvach et al. 2016), *iv)* long temporal coverage coupled with wide host taxonomic coverage within collections.

### 2.2. Parasite collection

We focus on gill parasites of haplochromines, because, contrary to many internal parasites, they display higher host-specificity (Poulin et al. 2011), they can be easily morphologically identified to species level, and they mostly have a direct life cycle (i.e. no intermediate hosts that may cloud the target host-parasite interaction). In Lake Victoria, haplochromine gill parasitofauna is dominated by monogenean flatworms (Monogenea) and copepod crustaceans (Copepoda), both with a direct life cycle. Monogeneans are hermaphrodite, with a free-swimming short-lived larval stage (Paperna 1996), and show high host specificity. Parasitic copepods have several larval stages, typically free-swimming, with only the adult female being a generalist parasite (Boxshall and Halsey 2004). In particular, the monogenean *Cichlidogyrus* is an emerging model for the evolution of host-parasite interactions (Cruz-Laufer et al. 2021, Vanhove et al. 2024).

We examined gill arches (right side of the fish only) under a dissecting stereoscope for gill macroparasites. The right side only was screened to minimize damage to collection specimens and to allow inter-studies comparability (as this is standard procedure, e.g. MacColl 2009, Gobbin et al. 2023). No differences have been detected in parasite abundance between the two gill sides (Roux et al. 2011). Formalin-fixed monogenean parasites (both recent and historical fish collections) were directly mounted on slides in Hoyer’s medium and examined with a microscope (Leica DM 2500 LED and Olympus BX41TF) under 1000x magnification. Ethanol-fixed monogenean specimens (recent fish collection only) were first cut into three parts: the two body extremities with sclerotized organs were mounted as described, the central body part was preserved in ethanol at 4°C for future molecular analysis. Monogenean species were identified following relevant literature (Vanhove et al. 2011, Gobbin et al. 2024) based on shape and size of sclerotized parts of the attachment organ (haptor) and of the male copulatory organ (MCO) (e.g. Paperna 1996, Grégoir et al. 2015, Gobbin et al. 2020, Gobbin et al. 2024), and counted. Other gill macroparasites were identified following Paperna (1996) and counted.

### 2.3. Statistical analysis

All statistical analyses were performed in R statistical software version 4.3.1 (R Core Team 2023). Since host species *Haplochromis nyererei* was disproportionately represented in the time period after perturbations (104 vs. 11 individuals), we randomly sampled 15 individuals from that time period (sample_n – dplyr package).

#### 2.4.1 Prevalence, intensity

Infection abundance incorporates both parasite **prevalence** (proportion of infected hosts) and **intensity** (number of parasites per infected host individual). We fit a Hurdle mixed model (glmmTMB – glmmTMB package; Magnusson et al. 2017) to simultaneously explore how parasite prevalence and intensity differed before vs. after perturbations and how it was influenced by parasite species. Hurdle models can handle zero-inflation, a common issue in parasitological analyses. Hurdle models have two components: a binary model fits data as zeroes or non-zeroes (prevalence) and, once this hurdle is crossed, a zero-truncated negative binomial model fits non-zero values (intensity, Zuur et al. 2009). Rank deficient parasite species were excluded from the hurdle model (i.e. *Gyrodactylus sturmbaueri*, cestode sp. I without data for one time period). The initial model included as fixed effects: time period (before vs. after perturbations), parasite species, and their interaction term. Initial random effects are: host species, fish individual identity (to account for individual variability in infection), fish length as an offset term (to account for the fact that bigger fish carry more parasites, Roux et al. 2011). Fish length was scaled prior to analysis (scale – base R, then adding 2 to avoid negative values) to balance fish interspecific differences in size. The optimal random effect structure was determined by Akaike information criterion (AIC) and Bayesian information criterion (BIC) comparisons (Schwarz 1978, Sakamoto et al. 1986). We used ANOVA (Anova – car package; Fox and Weisberg 2019) to estimate the parameters of significant fixed effects for both model components and thus select for the Minimum Adequate Model (MAM) (Nakagawa and Cuthill 2007). To compare parasite species before and after perturbations, we computed Estimated Marginal Means (emmeans – emmeans package) with custom contrasts and Benjamini-Hochberg correction for multiple comparisons (Benjamini and Hochberg 1995).

#### 2.4.2 Biodiversity metrics

We characterized gill parasite communities using parasite species richness per individual host (hereafter referred to as individual species richness) and **five biodiversity metrics** (so that each host individual is considered a living habitat for parasites): Shannon diversity index (accounting for parasite species richness and equitability; Shannon and Weaver 1949), Pielou’s evenness index (measuring the relative abundance of the different parasite species in a host community; Pielou 1977), Simpson’s diversity index (accounting for species richness and species relative abundance; Simpson 1949), Jaccard index (based on species presence/absence; Jaccard 1912), and Bray-Curtis distances (compositional dissimilarity based on parasite counts of infected hosts; Bray and Curtis 1957). We used Linear Mixed-Effects Models (lmer – lme4) to test whether the first four metrics changed after ecosystem perturbations. The initial model included time period (before vs. after perturbations), host species and their interaction term as fixed effects. Initial random effects were: host length and sampling effort per year (both as a regular random term or as an offset term). The optimal random effect structure and the MAM were determined as detailed above.

Since the Jaccard index and Bray-Curtis distances are expressed as a matrix, we used Analysis of Similarities (anosim – vegan) to test whether they changed after perturbations. We performed pairwise comparisons between host species in PAST3 (Hammer et al. 2001) (as this is currently unavailable in R), with Benjamini-Hochberg correction (p.adjust – base R).

#### 2.4.3 Host range and co-infections

The **host range** (number of host species infected by a given parasite species) was calculated per sampling year, excluding uninfected host individuals. Using Generalized Linear Models (glm – base R) we tested host range for the influence of: perturbations (before vs. after perturbations), parasite species, and their interaction. We also included an offset term, to account for the fish sampling effort of each year.

We tested whether **coinfections** (occurrence of two or more parasite species on the same host individual) were more frequent before or after ecosystem perturbations, using Generalized Linear Models (glmer.nb – lme4). Co-infections represent parasite-parasite interactions, an often-understudied aspect of parasitism that may be influenced by ecosystem changes. First, we calculated the number of parasite species per infected host individual and classified the infection as single- or multiple-species. The initial model included: time period (before vs. after perturbations), host species and their interaction term as fixed effects; fish identity as random effect, and host length as an offset (as described above). For both models, we followed the same procedure detailed above.

#### 2.4.4 Network analyses

We performed **network analyses** at host-individual level (infected host individuals only), rather than at species level, to consider the host intraspecific variation in parasite communities (e.g. Llopis-Belenguer et al. 2020, Llaberia-Robledillo et al. 2022). Indeed, a host individual often interacts with more than one parasite species (co-infections) and thus its role within the network varies depending on how many parasites it can transmit to other hosts (Runghen et al. 2021). For each network, before and after perturbations, we calculated three indices: *i)* connectance (C, Dunne et al. 2002), the proportion of realized links among all possible links (estimated with networklevel – bipartite package); *ii)* modularity (Q; Newman and Girvan 2004), measuring how well the network can be grouped into different modules (estimated with computeModules – bipartite package); and *iii)* weighted nestedness based on overlap and decreasing fill (WNODF; Almeida-Neto and Ulrich 2011), measuring the hierarchical organization of the community. We estimated host-parasite network stability as a combination of modularity and nestedness. In antagonistic networks (e.g. host-parasites), the network stability is enhanced by increasing nestedness and/or decreasing modularity (Thébault and Fontaine 2010).

Network size influences WNODF and Q values (Dunne et al. 2002), thus we generated 1000 null model replicates of the real matrices (nullmodel – bipartite package; Dormann et al. 2009) using the swap.web algorithm to standardize initial estimates of these two indices. Then, we used bootstrap replicates by host individuals to generate 1000 random networks of each original matrix. The 95% confidence intervals for each index were calculated (following Llopis-Belenguer et al. 2020). We tested differences of the three indices (C, Q, WNODF) between the period before and after perturbations using non-parametric Mann–Whitney–Wilcoxon (wilcox.test – base R).

## 3. Results

We found 654 monogeneans belonging to seven species (*Cichlidogyrus bifurcatus*, *C. furu*, *C. longipenis*, *C. nyanza*, *C. pseudodossoui*, *C. vetusmolendarius*, *Gyrodactylus sturmbaueri*), 691 copepods belonging to two species (*Lamproglena monodi*, *Ergasilus lamellifer*), and 15 cestodes (larval stages) belonging to an unidentified taxon (cestode sp. I). Eight parasite species were shared among the two time periods, whereas *G. sturmbaueri* was observed only once after perturbations and the larval cestode sp. I was observed only before perturbations. Parasites unidentified to species level (because of damaged specimens, 8.7% of monogeneans, 3.3% of copepods) were included in quantitative (e.g. prevalence, intensity) but not in qualitative analyses (e.g. network analyses), because there is no reason to assume these represent additional different species.

### 3.1. Prevalence and intensity

Infection **prevalence** decreased from 88.73% to 69.70% after perturbations in Lake Victoria (ChiSq_1_=28.747, p<0.001, **Fig 1**). The decrease in prevalence tended to vary across parasite species, although this was probably due to one parasite species (*C. longipenis*) deviating from the overall pattern (**Fig 2A**, **S1 Table**). Prevalence differed between parasite species (pooling time periods, ChiSq_9_=279.781, p<0.001): *Ergasilus lamellifer* had the highest infection prevalence (38.23%), whereas *C. bifurcatus* had the lowest (2.06%, **S1 Table**).

**Figure 1 -.**
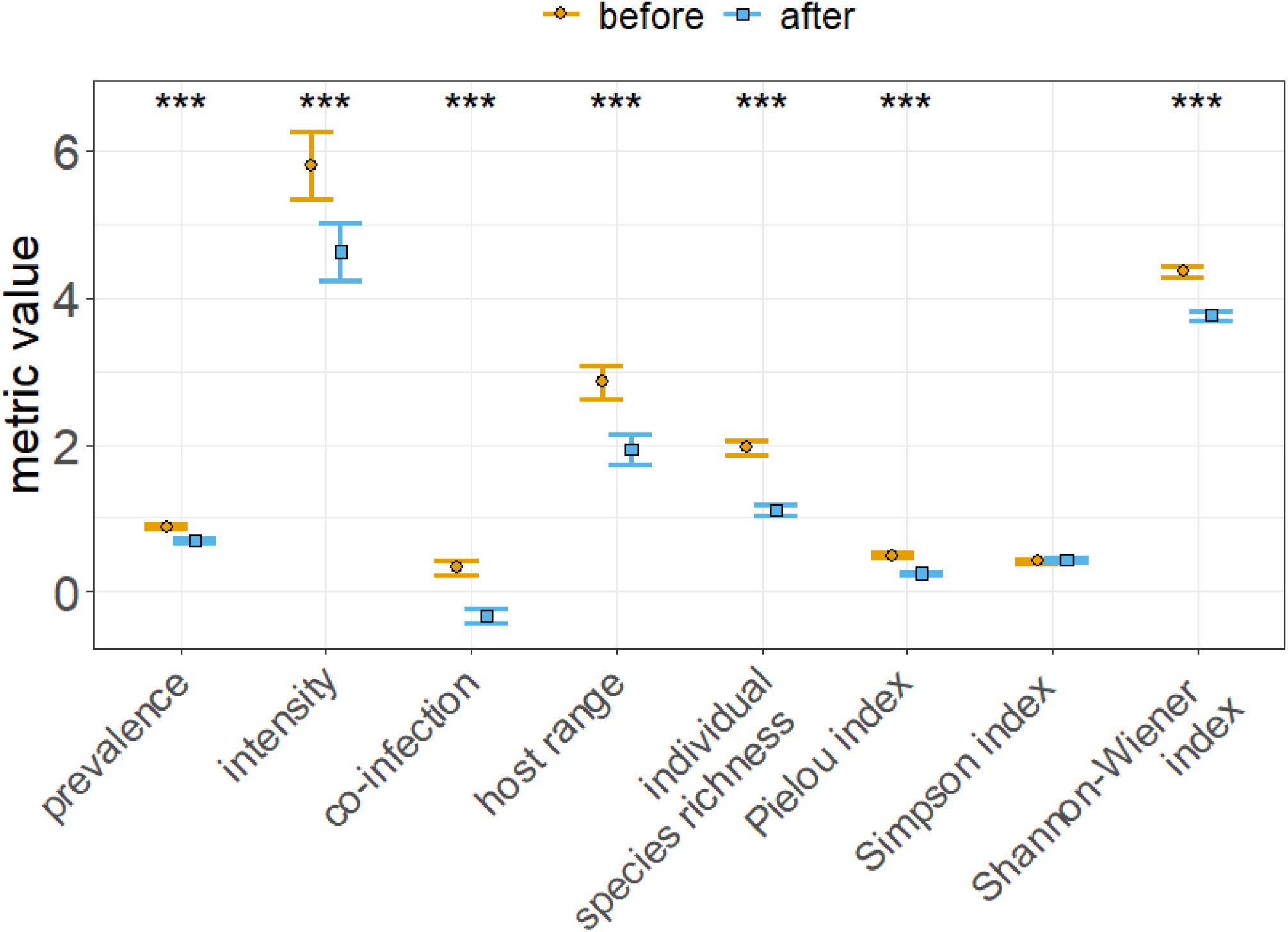
Infection prevalence, intensity, occurrence of co-infections and host range (number of host species exploited by a given parasite species), diversity metrics (individual species richness, Pielou’s evenness index, Shannon-Wiener index) decreased after perturbations in Lake Victoria. As an exception, the Simpson index did not change. Orange points: before perturbations, blue squares: after perturbations. ±SE

**Figure 2 -.**
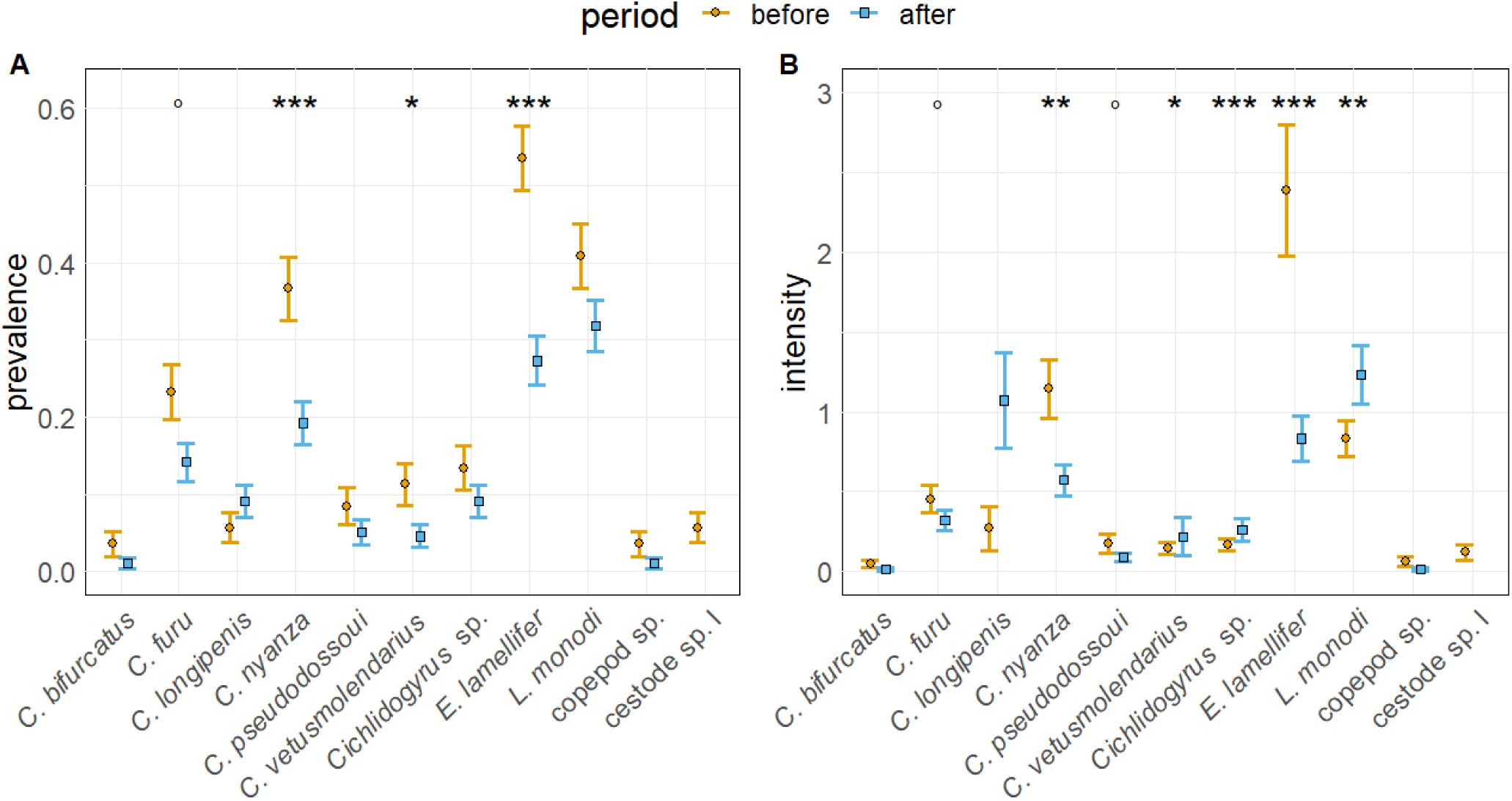
Infection prevalence **(A)** and intensity **(B)** ±SE were higher before (orange points) than after (blue squares) perturbations in Lake Victoria, overall and for some parasite species (significance level ° 0.1, * 0.05, ** 0.01 *** 0.001). Only one parasite species (*Cichlidogyrus longipenis*) increased in prevalence, and two others (*Cichlidogyrus vetusmolendarius, Lamproglena monodi*) increased in intensity.

Infection **intensity** decreased from 5.80 to 4.63 (ChiSq_1_=46.870, p<0.001, **Fig 1**) and this decrease varied across parasite species (ChiSq_9_=49.506, p<0.001). After ecosystem perturbations, three parasite species decreased or tended to decrease (*E. lamellifer* z=3.557, p<0.001; *C. nyanza* z=1.909, p=0.056; *C. pseudodossoui* z=1.816, p=0.069), whereas two parasite species increased in intensity (*C. vetusmolendarius* z=-2.871, p=0.004; *L. monodi* z=-2.912, p=0.0043; **Fig 2B**). Intensity also differed between parasite species (ChiSq_14_=830945, p<0.001): *C. longipenis* had the highest infection intensity (7.00), whereas *G. sturmbaueri* and *C. bifurcatus* had the lowest (1.00, 1.14, respectively, **S1 Table**).

### 3.2. Biodiversity indices

The individual species richness (Chisq_1_=47.312, p<0.001), the Shannon-Wiener index (Chisq_1_=38.100, p<0.001) and Pielou’s evenness index (Chisq_1_=29.794, p<0.001) decreased after perturbations, whereas the Simpson index did not change (Chisq_1_=0.199, p=0.655, **Fig 1**). This decrease varied across host species (**S2 Table**). Out of 13 host species, seven showed a decrease in both individual species richness and Shannon-Wiener index, while six decreased in Pielou’s evenness index (**S3 Table**). Host species differed in these indices (**S2 Table**).

Gill parasite communities differed between time periods (Bray-Curtis R2=0.017, F_1_=6.899, p=0.001; Jaccard R2=0.015, F_1_=6.187, p=0.001, **Fig 3**). This temporal variation in parasite communities varied across host species (Bray-Curtis R2=0.415, F_24_=7.15, p=0.001; Jaccard R2=0.431, F_24_=7.620, p=0.001): it was significantly different in nine host species (**S4 Table**).

**Fig 3 -.**
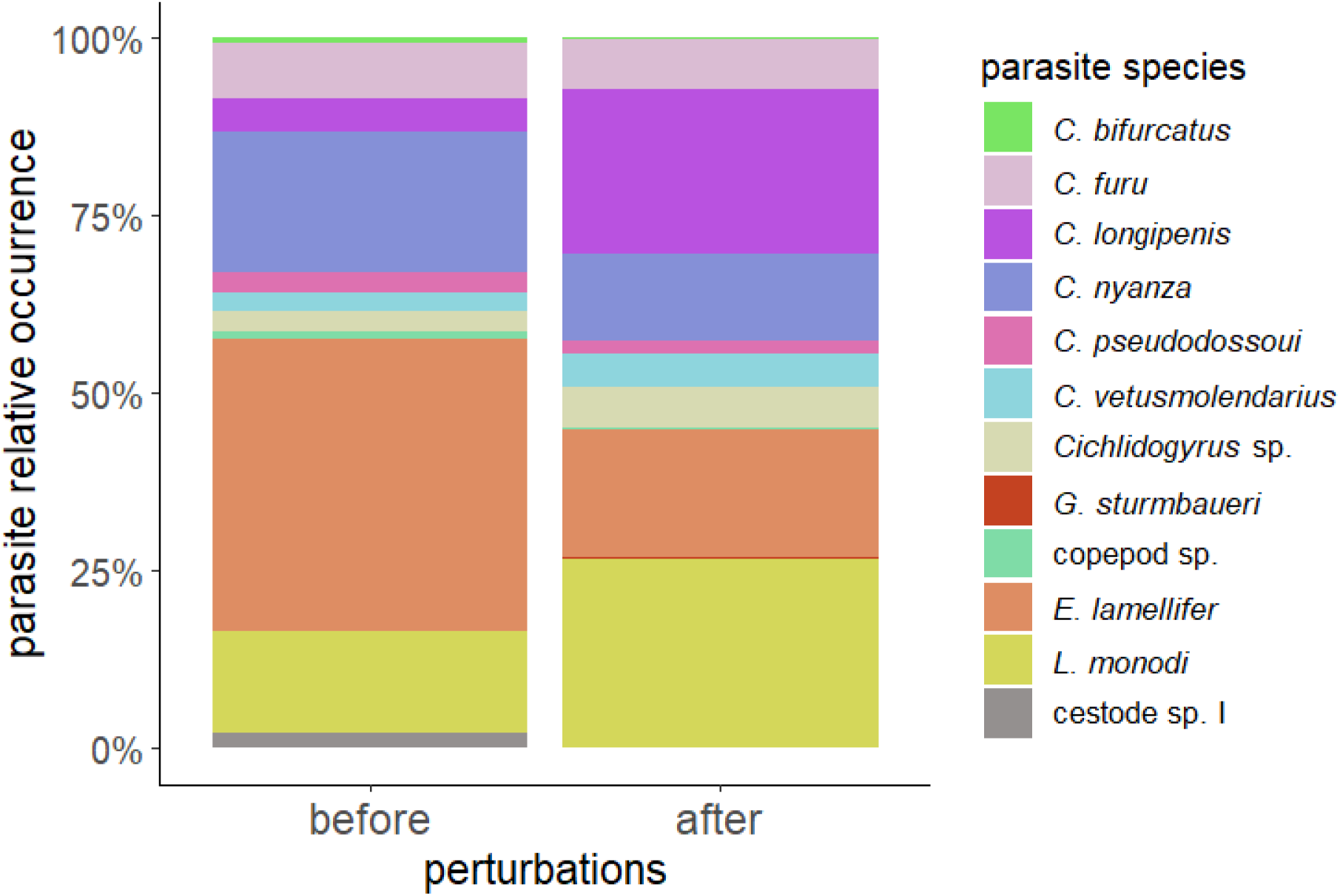
Parasite relative occurrence before and after perturbations in Lake Victoria. Most parasite species have been observed in both time periods, although their relative occurrence changed. Cestode sp. I was only observed before perturbations, whereas *Gyrodactylus sturmbaueri* was only observed after perturbations.

### 3.3. Host range and co-infection

The **host range** of parasites decreased significantly after perturbations (Chisq_1_=9.525, p=0.002, **Fig 1**), after accounting for the variation between parasite species (Chisq_11_=42.207, p<0.001). This decrease did not vary across parasite species (Chisq_9_=4.376, p=0.885). On average, the two copepod species had the largest host range (pooling time periods *L. monodi* 3.6, *E. lamellifer* 3.1 host species), whereas *C. longipenis* had the smallest (1 host species in this dataset, *A. alluaudi*).

Occurrence of co-infection decreased after ecosystem perturbations from 66.4% to 43.8% (Chisq_1_=13.771, p<0.001, **Fig 1**), after accounting for the variation between host species (Chisq_12_=25.927, p=0.011). Variation in parasite co-occurrence among host species did not differ between the two time periods (Chisq_12_=10.376, p=0.583).

### 3.4. Network analysis

The host-parasite networks differed between time periods (i.e. before and after perturbations) in their nestedness and modularity (both p<0.001), but they did not differ in their connectance (**Fig 4**). Standardized modularity was higher (0.51 vs. 0.35) and standardized nestedness was lower (14.16 vs. 24.56) after perturbations. Connectance was low in both periods (0.23 before and 0.18 after perturbations), which is due to low species richness of the network. After perturbations, some host-parasite combinations were no longer reported, while new ones were observed (**Fig 5**).

**Figure 4 -.**
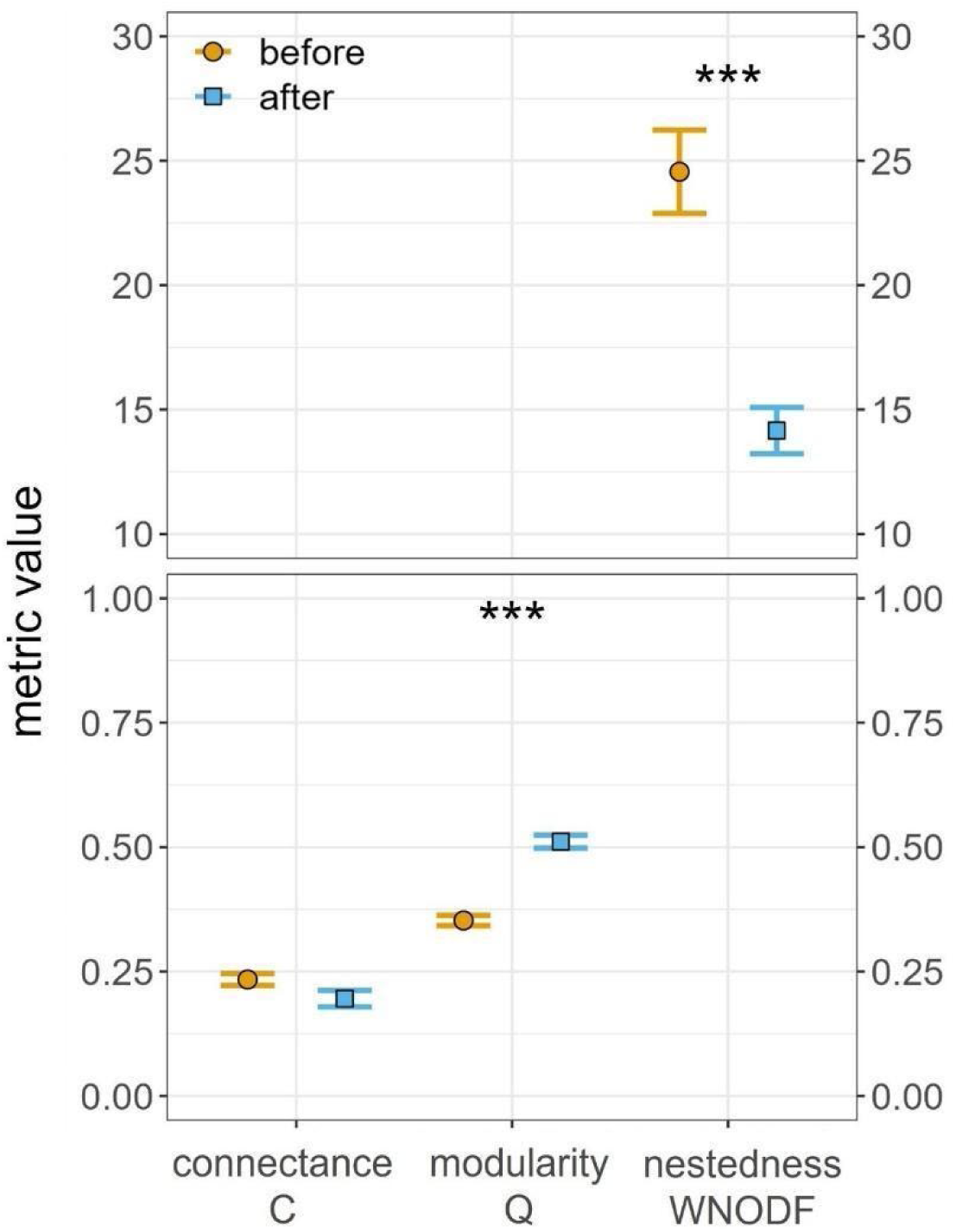
After perturbations, modularity (Q) increased whereas nestedness (WNODF) decreased in the host-gill parasite network. Connectance (C) did not change. Modularity and nestedness were standardized based on null.swap algorithm. Orange points: before perturbations, blue squares: after perturbations. ±SE.

**Fig 5 -.**
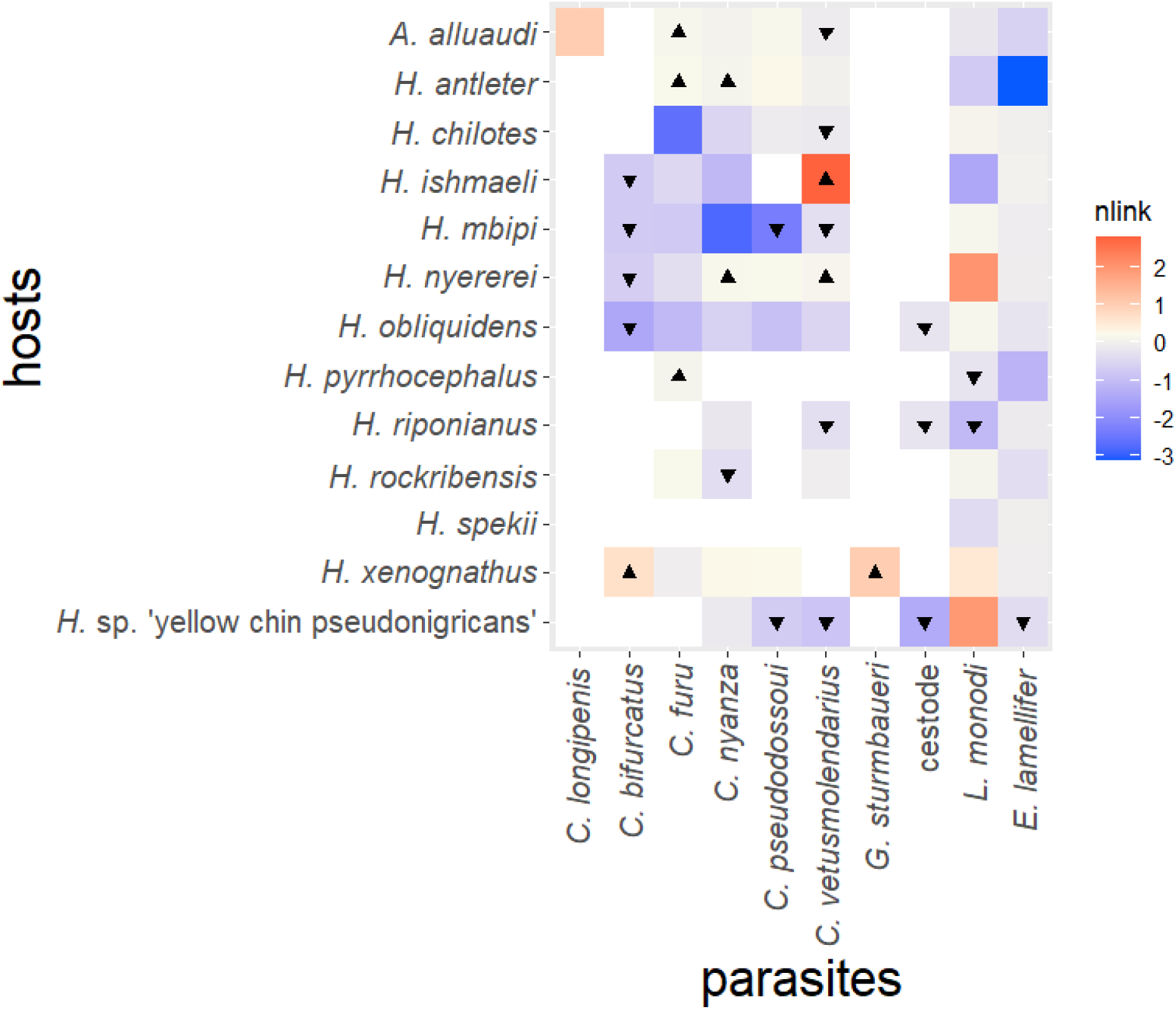
After perturbations in Lake Victoria, some host-parasite combinations increased in frequency (red) while other decreased (blue). New host-parasite combinations emerged after perturbations (▲), while others disappeared (▾). The darker the colour, the more times the parasite was found on that host species. White squares represent combinations neither observed before or after perturbations.

## 4. Discussion

We tested how parasites respond to anthropogenic perturbations in Lake Victoria, a biodiversity hotspot that experienced human-induced Nile perch invasion and eutrophication. We compared parasite communities before and after the onset of these perturbations, using recent and historical fish collections. Host-parasite interactions were affected by anthropogenic perturbations: the host-parasite network rearranged (**Fig 4, 5**) and infection parameters (intensity, prevalence, co-infections), host range, and biodiversity indices (individual species richness, Shannon-Wiener, Pielou’s evenness) decreased (**Fig 1, 2**). Since parasite co-introductions can be ruled out (see **Supplementary**), observed changes in parasitism are likely due to ecosystem changes.

Most parasite species showed a declining pattern in both prevalence and intensity, while these parameters increased for few others (**Fig 2**). This is in line with the “few winners, many losers” paradigm, which was already observed in other host-parasite systems (e.g. bat flies at different levels of anthropogenic disturbance, Pilosof et al. 2012). It is unclear why the winners are *L. monodi* and *C. vetusmolendarius*. We speculate it may be because *L. monodi* has a wide host range (19 genera of Cichlidae, but also one species of Alestidae, one of Clariidae, one of Cyprinidae; Ayawei 2023) combined with high prevalence and potentially high dispersal, whereas *C. vetusmolendarius* showed the ability to switch host species (present study). Prevalence and intensity changes were always mirrored by a change in the same direction in the relative occurrence of a given parasite within the parasite community (**Fig 3**), suggesting that parasite species declining in intensity and prevalence are also representing a smaller part of the infection; whereas parasites increasing in intensity (*C. vetusmolendarius*, *L. monodi*) are expanding into these emptied niches.

The aforementioned changes in infection parameters and in the host-parasite network occurred despite a consistent parasite species composition (i.e. same parasite species were observed before and after perturbations, except cestode sp. I and *Gyrodactylus sturmbaueri*, see **Supplementary** for details). This highlights that the absence of changes in species richness does not imply ecosystem stability, because other facets – such as host range, multispecies infection, bipartite network – could change through time. These aspects are ‘hidden’ because parasites are often simply numerically counted, without species-level identification. Therefore, it is important to monitor parasite communities and their interactions with their hosts to be able to detect ecosystem changes.

Lake Victoria is a young system; therefore, its host-parasite network is expected to be simple and relatively small, as we observed. Comparing *Cichlidogyrus*-cichlid networks in young Lake Victoria and ancient Lake Tanganyika (Cruz-Laufer et al. 2022) also supports the hypothesis that younger lakes have lower monogenean diversity (Pariselle et al. 2015). Host-parasite networks are usually modular and not nested (Fortuna et al. 2010, Thébault and Fontaine 2010, Morrison and Dirzo 2020). Disruption of the host-parasite network structure has already been observed after biological invasions (Runghen et al. 2021, Llaberia-Robledillo et al. 2022). Even if in Lake Victoria the parasite species richness mostly remained the same over time, the network rearranged: modularity increased and nestedness decreased after perturbations (**Fig 4**). These changes support our hypothesis of a network becoming less stable with anthropogenic changes, as stability of antagonistic networks is enhanced by increasing nestedness and/or decreasing modularity (Thébault and Fontaine 2010). The pattern we found is the opposite of those observed in a fish-parasite network after biological invasion (Llaberia-Robledillo et al. 2022) and in seed-dispersal networks after human impacts (Sebastián-González et al. 2015). However, caution is needed in comparing different network types, as they often differ in their characteristics (Takemoto and Iida 2019). An increase in **modularity** in response to anthropogenic perturbations was previously observed in host-parasite networks (Gilarranz et al. 2016, Lula Costa et al. 2023). The observed increase in modularity was not linked to an increase in infection intensity, as found by Lula Costa et al. (2023), suggesting that both changes resulted from perturbations rather than one being the effect of the other. The observed **nestedness** decrease is in contrast with the concept that a low nestedness is promoted when parasite niches are well established (Almeida-Neto and Ulrich 2011). Gill parasite taxa are shared among haplochromines of Lake Victoria, rather than being specialized to only one or few host species (Gobbin et al. 2020). Thus, these host-parasite interactions may be less vulnerable to perturbations than highly specialized interactions, making the network relatively resistant to perturbations considering its simplicity.

The ecological niche of parasites (i.e. host range) narrowed after perturbations (**Fig 1**), in line with a distribution range contraction driven by human activities observed in many mammal taxa (Laliberte and Ripple 2004, Pacifici et al. 2020). The current narrow niche likely results from a loss of host availability (both in terms of abundance and species richness), rather than from a real specialization process of the parasite. The ‘forced’ narrow niche poses an increased conservation risk: if more host species disappear, parasites may not be able to maintain a viable population. Thus, a niche contraction implies a decrease in resilience (i.e. a decreased ability to recover after perturbations). With many parasite species narrowing their niches, we may infer a general decrease in the resilience of the ecosystem of Lake Victoria. Parasites in Lake Victoria experiencing a contracting host range became more vulnerable and require at least monitoring of their conservation status. We highlight that a host range shift differs from a host shift: the former implies a quantitative change in host species exploited (decrease/increase in numbers) whereas the latter focuses on qualitative changes in host species exploited (colonization of different host species). While host shifts are often considered in disease literature (zoonosis and spillovers), changes in host range are overlooked. Along with host range contraptions, we observed host shifts: after perturbations some parasites no longer infected certain host species, while some parasites infected new host species (**Fig 5**). We highlight here the importance of monitoring changes in host range for early detection of ecosystem changes.

Parasitic infections are often studied in isolation, whereas in nature different parasite species commonly co-occur in the same host individual (Petney and Andrews 1998). Co-infections can have evolutionary (e.g. on virulence, Rigaud et al. 2010) and ecological consequences (e.g. on competition and niche distribution, Gobbin et al. 2021). Conversely, co-infections might be influenced by ecological changes and thus they need to be monitored in the context of global change. We observed a decrease in the occurrence of co-infections after perturbations (**Fig 1**). This may be linked to the decrease in parasite prevalence and intensity, which leads to a decrease in chances of multi-infection. The decrease of co-infections may also be due to *i)* parasites becoming less competitive and thus less able to infect host individuals in which another parasite species occurs and/or *ii)* a decreased facilitating effect (i.e. infection subsequent to mechanical damage, Bandilla et al. 2006; or through immunosuppression of the host, Zhi et al. 2018). Competition has been observed among species of *Cichlidogyrus* and hypothesized to have caused their niche distribution within host gills (Gobbin et al. 2021). The few parasite species that increased in intensity (*C. vetusmolendarius*, *L. monodi*) were not more likely to outcompete other parasite species, because their probabilities to occur in co-infections were not higher than those of parasites declining in intensity.

Parasite populations are declining in haplochromines of Lake Victoria and in other hosts elsewhere (Russell et al. 2015, MacKenzie and Pert 2018, Wood et al. 2023b). However, this trend cannot be generalized, as the direction of change (if any) can vary between parasite taxa even within the same study (Howard et al. 2019, Fiorenza et al. 2020b, Quinn et al. 2021, Welicky et al. 2021). A decline in abundance (which is an infection parameter incorporating both prevalence and intensity) is mostly observed in ectoparasites and in parasites with more than one host than endoparasites and parasites with one host (Carlson et al. 2017, Vanhove et al. 2022, Wood et al. 2023b). It is therefore important to monitor each ecosystem, as different parasites can respond differently in different ecosystems and to different perturbations. The decline in parasitism in Lake Victoria is detectable already in a 40-years period (20 years after the onset of perturbations). This 40-years’ timeframe is similar to that of previous studies in fish parasites (90 years in Welicky et al. 2021; 50 years in Fiorenza et al. 2020b). Further studies are needed to assess at higher temporal resolution how long it actually took for these changes in parasite communities to manifest.

The overall decline in parasitism in Lake Victoria and elsewhere, and the potential loss of one parasite taxon (cestode sp. I) stresses the urgent need for a conservation plan of parasites infecting wildlife (as already highlighted by Carlson et al. 2020). In response to this need, two of the authors recently established, together with other parasitologists worldwide, the IUCN Species Survival Commission’s Parasite Specialist Group, aiming at assessing the conservation status of parasite species globally (Hopkins and Kwak 2023).

## 5. Conclusion

In a relatively short period, human-induced perturbations in Lake Victoria affected not only free-living organisms, but also their parasites. Although gill parasite community studied here mostly maintained the same species, we observed a change in parasite community composition and an overall decline in infection and biodiversity metrics. The decrease in infection intensity, prevalence and host range poses a conservation risk for parasites. The host-parasite network re-arranged in a way that indicates ecosystem instability (i.e. modularity increased, nestedness decreased). The host range of parasites (which is linked to resilience ability) decreased and thus we expect a decreased resilience in the future. We showed the complementarity of two approaches, bipartite network and infection parameters, in detecting temporal variation in host-parasite interactions. Parasites contribute to biodiversity and increase enormously the number of connections between organisms, which is crucial for ecosystem resilience (especially in the face of current global change). It is therefore important to monitor these metrics of parasitism across time, rather than merely assess species richness, to evaluate both the conservation status of parasite species and the health status of ecosystems, and subsequently enforce conservation policies.

