## Supplementary for "Worms of change: anthropogenic disturbance changed the ectoparasite community structure of Lake Victoria cichlids"

page 1	Ruling out parasite co-introductions

Considerations about the parasite species observed in one time period only

page 2	Table S1

Table S2

page 3	Table S3

Table S4

page 4	References

**Ruling out parasite co-introductions**

Gill parasites have been co-introduced into Lake Victoria with two species of tilapia (*Oreochromis niloticus*, *O. leucostictus*) and Nile perch *(Lates niloticus).* Only two gill parasites of tilapias have been observed on conspecific hosts in Lake Victoria: the monogenean *Scutogyrus longicornis*, which was never observed on African halpochromines; Scholz et al. 2018), and the copepod *Lamproglena monodi*, which is a generalist found on 22 fish genera of 3 families, including haplochromines in Lake Victoria (Ayawei 2023). Among Nile perch parasites, no monogeneans have been found on any African freshwater fish other than species of *Lates* (Scholz et al. 2018, Kmentová et al. 2020). Three crustaceans species infecting Nile perch also infect haplochromines elsewhere, but were never reported in Lake Victoria (not on *L. niloticus* nor on haplochromines, Outa et al. 2021). Unsurprisingly, gill parasites of the Nile perch and of tilapias did not switch to haplochromines in Lake Victoria, indicating that changes observed in parasite communities of Victorian cichlids are not due to parasite (co-)introductions.

**Considerations about the parasite species observed in one time period only**

The one individual of *Gyrodactylus* recorded after perturbations may be an accidental infection, or it may represent the low extremity of these worms’ wide variability in prevalence in African cichlids (Zahradníčková et al. 2016, Jorissen et al. 2020). Cestode sp. I was observed only before perturbations. Cestodes have indirect life cycles (i.e. more than one host), thus they face a higher conservation risk when a step of the cycle is disrupted (e.g. if a host goes extinct or its population declines; Rohr et al. 2011) and are more dependent on host trophic connections than parasites with direct life cycles. The decline in host diversity and abundance that followed perturbations in Lake Victoria may have led to a decline of this cestode taxon. Cestodes generally have low host specificity, making them prone to switch host species. It is therefore possible that it switched to a host species not surveyed in this study. Since the cestode larvae were not identified to finer resolution, we can depict two alternative scenarios: *i)* the taxon is endemic to Lake Victoria, thus it may be threatened or extinct (if it does not infect or did not switch to other hosts), *ii)* the taxon is not endemic to Lake Victoria and persists in other host species and other lakes of the region. Either way, we can speculate that the Lake Victoria populations of cestode sp. I have been heavily impacted by perturbations and/or that it had a lower resilience than other parasite species (as parasites with indirect life cycles are less resilient, Wood et al. 2023).

**Table S1** - Infection prevalence and intensity overall and for each parasite species, before and after perturbations in Lake Victoria. *Cichlidogyrus* spp. are specimens of *Cichlidogyrus* that were not suitable for identification to species level, but there is no reason to assume these represent additional different species. We switched the sign of estimates of the binary model (prevalence) so that negative values represent a decrease in the probability of fish being infected.

χ2: Chi square for overall comparison, z: z-ratio for parasite species comparisons.

|  | **Prevalence** | | | | | |  | **Intensity** | | | | | |
| --- | --- | --- | --- | --- | --- | --- | --- | --- | --- | --- | --- | --- | --- |
|  | **tot** | **before** | **after** | **χ2 z** | **p** |  |  | **tot** | **before** | **after** | **χ2 z** | **p** |  |
| overall | 0.78 | 0.89 | 0.70 | 31.317 | **<0.001** | *** |  | 4.03 | 5.80 | 4.63 | 42.87 | **<0.001** | *** |
| *Cichlidogyrus bifurcatus* | 0.02 | 0.04 | 0.01 | -1.489 | 0.137 |  |  | 0.02 | 0.05 | 0.01 |  |  |  |
| *Cichlidogyrus furu* | 0.18 | 0.23 | 0.14 | -1.932 | **0.053** | . |  | 0.30 | 0.45 | 0.32 | 1.32 | 0.188 |  |
| *Cichlidogyrus longipenis* | 0.08 | 0.06 | 0.09 | 1.226 | 0.220 |  |  | 0.54 | 0.27 | 1.07 | -1.16 | 0.245 |  |
| *Cichlidogyrus nyanza* | 0.26 | 0.37 | 0.19 | -3.504 | **0.001** | *** |  | 0.66 | 1.14 | 0.57 | 1.91 | **0.056** | . |
| *Cichlidogyrus pseudodossoui* | 0.06 | 0.08 | 0.05 | -1.203 | 0.229 |  |  | 0.10 | 0.17 | 0.09 | 1.82 | **0.069** | . |
| *Cichlidogyrus vetusmolendarius* | 0.07 | 0.11 | 0.05 | -2.234 | **0.026** | * |  | 0.14 | 0.14 | 0.22 | -2.87 | **0.004** | ** |
| *Cichlidogyrus* spp. | 0.11 | 0.13 | 0.09 | -1.194 | 0.233 |  |  | 0.17 | 0.17 | 0.26 | -2.45 | **0.014** | * |
| copepod spp. | 0.04 | 0.08 | 0.01 | -1.489 | 0.137 |  |  | 0.06 | 0.15 | 0.01 |  |  |  |
| *Ergasilus lamellifer* | 0.38 | 0.53 | 0.27 | -4.691 | **<0.001** | *** |  | 1.22 | 2.38 | 0.83 | 3.56 | **<0.001** | *** |
| *Gyrodactylus sturmbaueri* | 0.00 | 0.00 | 0.01 |  |  |  |  | 0.00 | 0.00 | 0.01 |  |  |  |
| *Lamproglena monodi* | 0.36 | 0.41 | 0.32 | -1.637 | 0.102 |  |  | 0.78 | 0.75 | 1.23 | -2.91 | **0.004** | ** |
| cestode sp. I | 0.02 | 0.06 | 0.00 |  |  |  |  | 0.04 | 0.12 | 0.00 |  |  |  |

**Table S2** - Infection diversity measures (individual species richness, Pielou’s evenness index, Shannon-Wiener) index overall and for each host species. Simpson index not shown because it did not change after perturbations.

| **Model terms tested** | **Pielou's evenness index** | |  | **Individual species richness** | |  | **Shannon-Wiener index** | |  |
| --- | --- | --- | --- | --- | --- | --- | --- | --- | --- |
|  | **Chisq** | **p** |  | **Chisq** | **p** |  | **Chisq** | **p** |  |
| difference between host species | 48.831 | <0.001 | *** | 76.445 | <0.001 | *** | 65.03 | <0.001 | *** |
| difference between time periods | 29.794 | <0.001 | *** | 47.312 | <0.001 | *** | 38.10 | <0.001 | *** |
| temporal variation between host species | 26.426 | 0.0093 | ** | 31.169 | 0.0019 | ** | 36.53 | <0.001 | *** |

**Table S3** - Infection diversity measures (individual species richness, Pielou’s evenness index, Shannon-Wiener) index overall and for each host species. Simpson index not shown because it did not change after perturbations. χ2: Chi square for overall comparison, z: z-ratio for host species comparisons.

| **Host species** | **Pielou's evenness index** | | |  | **Individual species richness** | | |  | **Shannon-Wiener index** | | |
| --- | --- | --- | --- | --- | --- | --- | --- | --- | --- | --- | --- |
| **before vs. after perturbations** | **χ2** | **p** |  |  | **χ2** | **p** |  |  | **χ2** | **p** |  |
|  | **z** |  |  |  | **z** |  |  |  | **z** |  |  |
| overall | 29.794 | **<0.001** | *** |  | 47.312 | **<0.001** | *** |  | 38.100 | **<0.001** | *** |
| *A. alluaudi* | 0.989 | 0.323 |  |  | 1.792 | 0.074 | . |  | 1.126 | 0.261 |  |
| *H. chilotes* | 3.886 | **<0.001** | *** |  | 4.874 | **<0.001** | *** |  | 4.571 | **<0.001** | *** |
| *H. sauvagei* | 2.873 | **0.004** | ** |  | 2.237 | **0.026** | * |  | 2.436 | **0.015** | * |
| *H.* sp. 'yellow chin pseudonigricans' | 2.579 | **0.01** | ** |  | 3.190 | **0.002** | ** |  | 3.266 | **0.001** | *** |
| *H. xenognathus* | -0.891 | 0.374 |  |  | -0.834 | 0.405 |  |  | -0.983 | 0.327 |  |
| *H. antleter* | -0.230 | 0.819 |  |  | 0.111 | 0.912 |  |  | -0.058 | 0.954 |  |
| *H. ishmaeli* | 2.677 | **0.008** | ** |  | 2.358 | **0.019** | * |  | 2.647 | **0.009** | ** |
| *H. mbipi* | 2.436 | **0.016** | * |  | 3.768 | **<0.001** | *** |  | 2.973 | **0.003** | ** |
| *H. nyererei* | 0.005 | 0.996 |  |  | -0.227 | 0.821 |  |  | -0.279 | 0.780 |  |
| *H. obliquidens* | 2.658 | **0.008** | ** |  | 2.807 | **0.005** | ** |  | 3.641 | **<0.001** | *** |
| *H. pyrrhocephalus* | 0.864 | 0.388 |  |  | 2.544 | **0.011** | * |  | 0.611 | 0.542 |  |
| *H. riponianus* | 1.513 | 0.131 |  |  | 1.287 | 0.199 |  |  | 1.991 | **0.047** | * |
| *H. spekii* | -0.504 | 0.614 |  |  | 0.278 | 0.781 |  |  | -0.356 | 0.722 |  |

**Table S4** - Comparison of Jaccard index and Bray-Curtis distances before and after perturbations within host species. Bray-Curtis distances differed in nine host species, whereas Jaccard index differed in three host species.

| **Host species** | **Jaccard** | | **Bray-Curtis** | |
| --- | --- | --- | --- | --- |
| *A. alluaudi* | **0.007** | ** | **0.001** | *** |
| *H. antleter* | 0.073 | . | **0.008** | ** |
| *H. chilotes* | 0.170 |  | **0.038** | * |
| *H. ishmaeli* | 0.321 |  | **0.008** | ** |
| *H. mbipi* | 0.393 |  | **0.038** | * |
| *H. nyererei* | **0.007** | ** | **0.006** | ** |
| *H. obliquidens* | 0.340 |  | 0.139 |  |
| *H. riponianus* | 0.393 |  | 0.337 |  |
| *H. sauvagei* | **0.001** | *** | **0.006** | ** |
| *H.* sp. 'yellow chin pseudonigricans' | 0.053 | . | **0.006** | ** |
| *H. spekii* | 0.393 |  | 0.337 |  |
| *H. xenognathus* | 0.415 |  | 0.714 |  |
| *H. pyrrhocephalus* | 0.393 |  | **0.013** | * |
